# Benchmarking Universal Machine Learning Force Fields with Hydrogen-Bonding Cooperativity

**DOI:** 10.1101/2025.04.29.651212

**Authors:** Xinping Feng, You Xu, Jing Huang

**Affiliations:** School of Life Sciences, Westlake University, Hangzhou, Zhejiang 310030, China; Westlake AI Therapeutics Laboratory, Westlake Laboratory of Life Sciences and Biomedicine, Hangzhou, Zhejiang 310024, China; Institute of Biology, Westlake Institute for Advanced Study, Hangzhou, Zhejiang 310024, China

**Keywords:** Machine learning force field, cooperative effects, self-assembly, neural network potential, hydrogen bond

## Abstract

Machine learning force fields (MLFFs) offer a promising balance between quantum mechanical (QM) accuracy and molecular mechanics efficiency. While MLFFs have shown strong performance in modeling short-range interactions and reproducing potential energy surfaces, their ability to capture long-range cooperative effects remains underexplored. In this study, we assess the ability of three MLFF models — ANI, MACE-OFF, and Orb — to reproduce cooperative interactions arising from environmental induction and dispersion, which are essential for many biomolecular processes. Using a recently proposed framework, we quantify hydrogen bond (H-bond) cooperativity in N-methylacetamide polymers. Our results show that all MLFFs capture cooperativity to some extent, with MACE-OFF yielding the closest agreement with QM data. These findings highlight the importance of evaluating many-body effects in MLFFs and suggest that H-bond cooperativity can serve as a useful benchmark for improving their physical fidelity.

## 1 Introduction

Machine learning potentials (MLPs), sometimes also referred to as machine learning force fields (MLFFs), have emerged as a significant advancement in computational chemistry, offering a balance between the high accuracy of quantum mechanics (QM) and the computational efficiency of molecular mechanics (MM) [1, 2]. By leveraging machine learning algorithms, MLPs learn the statistical relationship between molecular structures and their potential energies from large datasets. Various methodologies, including kernel-based approaches and neural network (NN) models, have demonstrated notable success in simulating and predicting the properties of complex chemical systems.

NN potentials are particularly promising because deep NNs excel at fitting high dimensional data distributions, enabling them to capture intricate intra- and intermolecular interactions [3, 4, 5, 6, 7, 8]. The seminal work of Behler and Parrinello introduced a scheme in which the total potential energy is decomposed into atomic contributions, each predicted by an NN that takes atomcentered environment descriptors as input [9]. The field has progressed rapidly since then, with more advanced network architectures designed to better preserve physical symmetries and improve training efficiency [10, 11, 12, 13, 14]. Whereas early MLPs were usually trained bespoke on data generated for the particular system studied, recent efforts have shifted toward pretraining models for broad, offtheshelf use — much like traditional force fields (FFs) that are carefully parametrized once and then distributed to endusers. Resonating with the wider ML move toward foundation models, an increasing number of universal MLFFs covering substantial regions of chemical space are now becoming available.

One of the most popular universal MLFFs is the ANI series developed by Roitberg and co-workers. The ANI-2x model, employs transfer learning: a network first trained on a large DFT dataset is finetuned with a smaller, highlevel CCSD(T) dataset, achieving accuracy across a broad chemical space that includes C, H, N, O, S, F, and Cl [15], and performing well on a wide range of drug-like molecules and small peptides. Another notable model, MACE, extends message-passing neural networks (MPNNs) by incorporating higher-order equivariant messages, boosting both efficiency and accuracy [13]. The MACE open force fields (MACE-OFFs) were trained on diverse datasets including organic molecules, water clusters, small peptides and dipeptides, and have demonstrated state-of-the-art performance not only in reproducing potential energy surfaces (PES) and atomic forces, but also in predicting condensed-phase properties such as liquid densities and solvation free energies [16, 17]. Meanwhile, the Orb model, designed for large-scalesimulations of inorganic and crystalline materials, employs a scalable graph NN architecture that preserves rotational invariance and allows efficient modeling of long-range dispersion interactions via diffusion pre-training and D3 correction [14].

In classical FFs, non-bonded interactions are evaluated throughout the space. In particular, electrostatics are typically handled with the particle meshed Ewald (PME) method, and more recently the LJ-PME method has been adopted to fully account for the van der Waals interactions[18, 19]. This contrasts with MLFFs, in which atomic energies are computed from the information contained within a direct cutoff (typically 4.0 Å to 6.0 Å). Interactions beyond this range are presumed to be captured indirectly, for example through multiple passes of message passing, yet the extent to which MLFFs reproduce full long range interaction remains to be systematically benchmarked [20]. Long-range interactions are essential in chemical and biological systems dominated by non-covalent forces, for example proteins. A prime example is the cooperative effects of hydrogen bonds (H-bonds) in stabilizing protein structures. Once a few H-bonds form between residues, subsequent H-bond formation becomes energetically more favorable, facilitating protein folding, assembly, and aggregation [21, 22, 23].

H-bond cooperativity arises mainly from electronic induction, reflecting the molecular polarizability in response to the surrounding environment. QM methods such as CCSD(T), MP2, and DFT can accurately capture these effects, as can polarizable MM force fields, which explicitly account for charge redistribution [24, 25, 26, 27]. In contrast, additive force fields fail to reproduce such cooperative effects. The quantification of cooperative energy has historically varied due to differences in computational methods, model systems, and definitions of cooperativity. QM calculations of various H-bonded polymers, including N-methylacetamide (NMA) [28], water [29], N-methylformamide (NMF) [30], formamide [31], and alanine peptides [32, 33], have yielded cooperative energy estimates ranging from 3 to 26 kcal/mol, depending on the specific formulation used.

Recent methodological advancements have enabled more rigorous quantification of cooperative effects and improved benchmark for classical MM force fields [34]. In this recent study, we introduced a general framework that defines cooperativity as the difference between the interaction energy of an isolated dimer 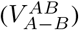 and its interaction energy in the presence of a third molecule 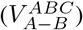 (Figure 1a). This cooperative energy is computed using the internal energies of optimized geometries (Equation 1).

**Figure 1.**
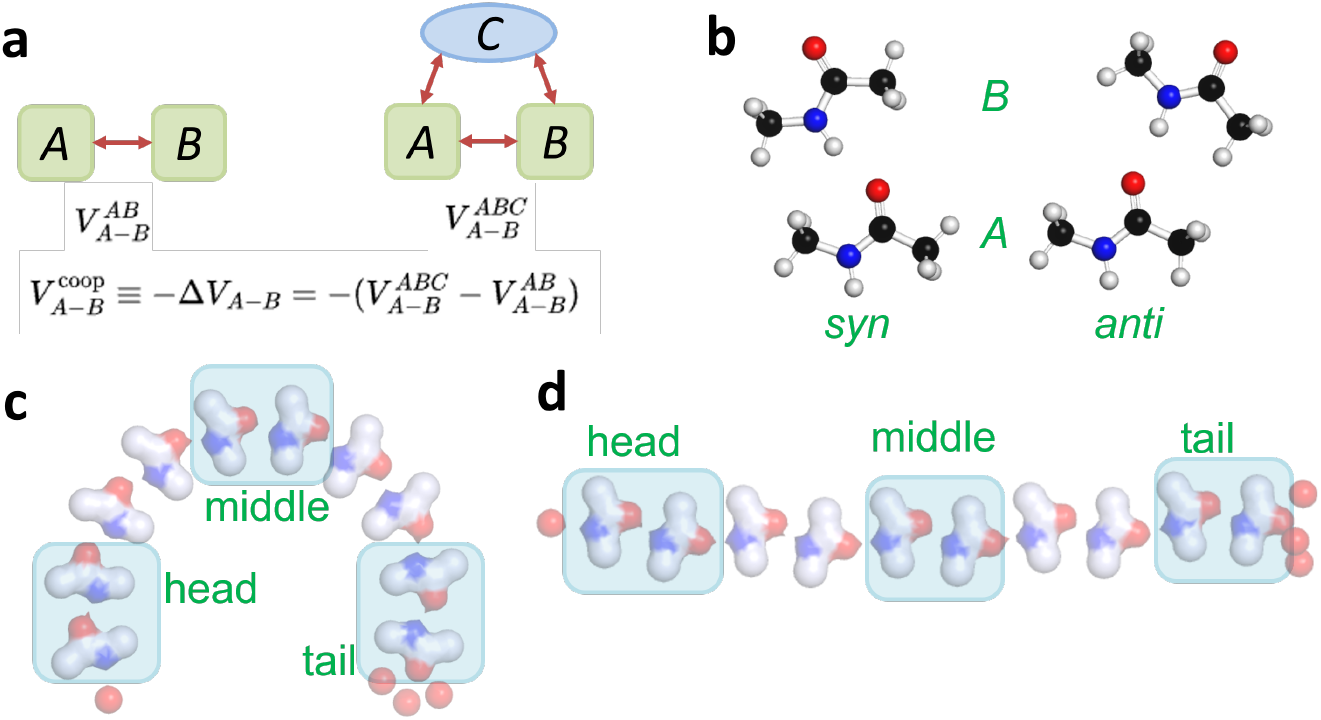
The scheme of cooperativity calculation and models. (a) Cooperativity is the difference of dimer interaction between two-body and three-body systems. (b) Two conformations of NMA dimer optimized at the *ω*B97XD/cc-pVTZ level of QM. (c) An arc decamer with exclusively syn conformation of dimer blocks. (d) A linear decamer with alternating syn and anti blocks. In (c) and (d), the water capping sites on termini are illustrated; and the cooperative energies calculated for the first, middle, and last H-bonds to evaluate the effect of NMAs extending on B-side, both sides and A-side, respectively, are highlighted with transparent boxes.

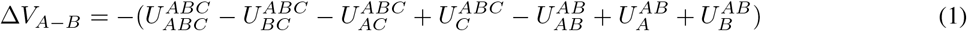

As the superscripts indicate, the first four terms (internal energies of *ABC, BC, AC*, and *C*) are computed using the geometry and basis set of complex *ABC*, while the remaining three terms (internal energies of *AB, A*, and *B*) are obtained using the geometry and basis set of complex *AB*. Here, *A, B*, and *C* represent any number of molecules. This framework accounts for the cooperative effects arising from both geometric distortion and electrostatic induction within molecules *A* and *B* in response to the addition of *C*, while explicitly excluding any contributions from the internal interactions or polarization of *C* itself. This formulation thus enables a systematic analysis of many-body effects in various molecular environments.

Here, we apply this theoretical framework to evaluate the ability of MLFFs to capture H-bond cooperatives in MLFFs. We used the same model systems and QM references, where NMA chains were optimized at the *ω*B97XD/cc-pVTZ level and energies computed at the RI-MP2/aug-cc-pVTZ level. Our findings reveal that all MLFFs considered, including the ANI, MACE-OFF, and Orb models, capture cooperativity to some extent but with considerable variance in accuracy. Despite being trained on high-quality QM data covering chemical spaces, these MLFFs differ substantially in how they represent environment-dependent interactions. Our work provides a critical benchmark for MLFFs and sheds light on how these models capture complex cooperative effects.

## 2 Method

Homogeneous NMA polymers identical to the previous study were used [34], where NMAs oriented in parallel configuration were arranged to form hydrogen bonds (H-bonds) in a head-to-tail way. As the fundamental H-bonded block, an NMA dimer was used, in which a hydrogen bond was formed between the carbonyl oxygen of molecule A (*O*_*A*_) and the amide nitrogen of molecule B (*N*_*B*_). The dimer was constrained to remain planar.

The NMA dimer adopted two distinct conformations. In the syn conformation, atoms *N*_*A*_ and *N*_*B*_ are positioned on the same side of the C_*A*_-O_*A*_ axis, corresponding to dihedral *ϕ*_1_ (N_*A*_-C_*A*_-O_*A*_-N_*B*_) of 0^*°*^. In contrast, in the anti conformation, *N*_*A*_ and *N*_*B*_ lay on opposite sides of the C_*A*_-O_*A*_ axis, with *ϕ*_1_ at 180^*°*^ (Figure 1b). Two well-ordered NMA polymer patterns were examined: one in which identical dimers formed an arc pattern and another where alternating syn and anti dimers resulted in a linear pattern. All polymer structures were built starting from a syn dimer and extended up to 12 NMAs. The head, middle, and tail H-bonds were calculated within each polymer to assess the cooperative effects of chain elongation on different sides (Figure 1c, d).

The cooperativity of H-bond interactions was evaluated for three families of MLFFs (Table 1), ANI models including 1ccx [35] and 2x [15], Orb models [14] including d3-v2, v2, and v3-conservative-20-omat [36], and MACE-OFF models including 23-S, 23-M, 23b-M [16], 23-SC [37], and 24-M [17]. The energy terms of equation 1 were calculated from the corresponding geometries that were energetically minimized. The initial conformations were taken from QM-optimized geometries. During minimization, torsions *ϕ*_1_ and *ϕ*_2_ (C_*A*_-O_*A*_-N_*B*_-C_*B*_) was constrained to maintain planarity across all NMA molecules. For Orb models, structure optimization using the L-BFGS algorithm and single-point potential energy calculations were performed via the ASE calculator interface [38]. For MACE-OFF and ANI models, structures were minimized using the L-BFGS algorithm, and potential energy calculations were carried out in OpenMM [39], employing a customized API (openmm-ml) designed for ML models in simulations (https://github.com/jharrymoore/openmm-ml/tree/main).

**Table 1:**
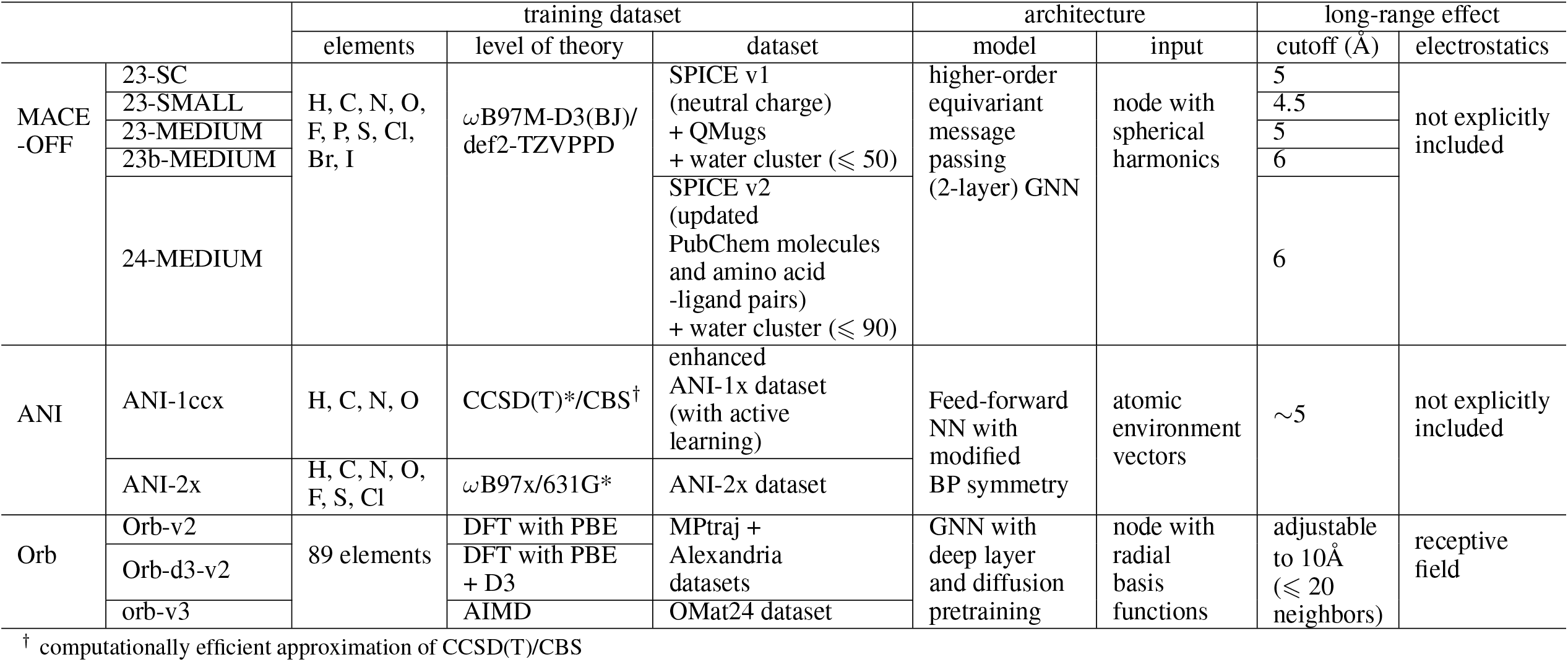
Feature comparison of the MLFFs evaluated in this study.

## 3 Results

### 3.1 NMA dimer interaction and conformation

NMA serves as a model system for protein backbone interactions in the parametrization of classical force fields. The H-bond interaction between the amide and the carbonyl group stabilizes NMA dimer in a planar conformation (Figure 1b, c). For longer NMA polymers, however, the global energy minimum usually corresponds to irregular aggregated structures with different potential energy models. To systematically evaluate the cooperative effects of each elongating NMA that mimic the formation of secondary structures, planarity restraints were applied to all polymers during optimization, including the reference dimer. In our previous QM calculations, both syn and anti conformations were optimized using density functional theory (DFT) at the *ω*B97XD/cc-pVTZ level, with interaction energies evaluated at the SCS-MP2/aug-cc-pVTZ level. These calculations indicated an energy difference of approximately 0.6 kcal/mol between the two conformations. Among MLFFs, all MACE-OFF models provided consistent interaction energy estimates, whereas ANI and Orb models exhibited greater variation across their respective variants (Figure 2). Most MLFF models overestimated the interaction energies, though the deviations for all models except ANI-2x and Orb-v2 remained within 2 kcal/mol, which is comparable to those observed in MM force fields.

**Figure 2.**
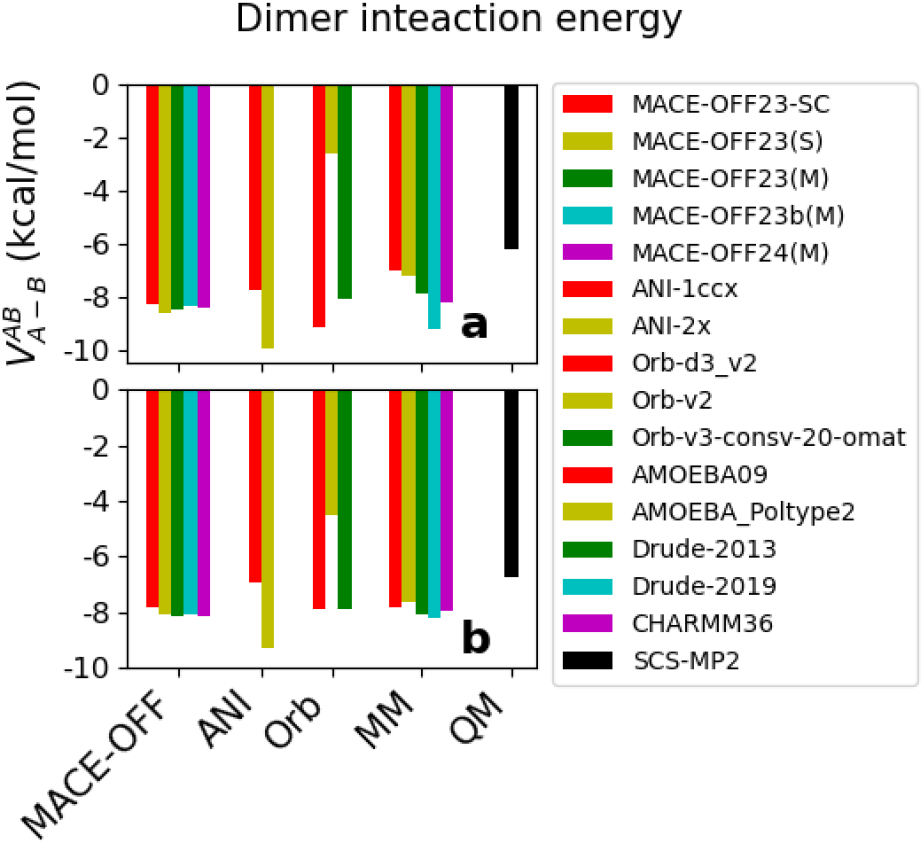
Dimer interaction of each model. (a) syn conformation; (b) anti conformation. The x-axis indicates the group of force fields and legend of variants is shown on the right.

Without restraints, geometric optimization using MACE-OFF and ANI still recovered the correct dimer conformations (Figure 1b, c), indicating that both force fields provide an adequate description of NMA interaction. In contrast, Orb-v2 models failed to reproduce the H-bonding conformation, instead favoring stacked structures (Figure S1a). This issue persisted for longer chains, where Orb-v2 consistently failed to achieve H-bonded conformations, even under planarity restraints (Figure S1b). To enforce a regular polymer pattern capable of capturing H-bond cooperativity, a much looser convergence tolerance (4.6 kcal/mol/Å) had to be applied on Orb-v2 compared to the 0.00024 kcal/mol/Å used for MACE-OFF and ANI. This limitation was partially addressed in the recently released Orb-v3-conservative-20-omat, which showed improved recovery of the H-bonded dimer and better polymer geometry optimization under reasonable convergence criteria. Nonetheless, Orb-v3 models continued to exhibit challenges when applied to long polymers in the presence of water. As a result, the cooperative energy for Orb models were calculated using QM geometries rather than Orb optimized polymer structures (Figure S1c).

### 3.2 Cooperative energy of H-bonds

Although the interaction energies of isolated dimers differ slightly between syn and anti conformations, the overall profile of cooperativity for arc and linear polymer patterns remains similar, as shown in the previous study. Such trend was also consistently observed across all MLFFs evaluated in this work. For clarity, the main text focuses on arc polymers composed of repeating syn dimers, while the corresponding results for linear polymers are provided in the Supporting Information (SI).

Cooperativity was present across all three MLFFs, confirming their reproduction in polarizability. However, the addition of NMAs to either side of the dimer produced markedly different cooperative effects among these models (Figure 3, Figure S2 for linear polymers). The MACE-OFF models closely reproduced the MP2 reference, whereas ANI models substantially underestimated and Orb models significantly overestimated cooperative energies. Consistent with the QM reference, the maximum cooperative energies on both terminal H-bonds were equivalent, measuring 1.4-1.6 kcal/mol in MACE variants, 0.3-0.6 kcal/mol in ANI, 7.5-10 kcal/mol in Orb-v2 models, and 3.5-4 kcal/mol in Orb-v3 model. The maximum cooperative energy for middle hydrogen bonds, as calculated by MP2, was 4.2 kcal/mol. This value was closely reproduced by MACE (2.8-3.2 kcal/mol), underestimated by ANI (0.4-1.0 kcal/mol), significantly overestimated by Orb-v2 (18-20 kcal/mol), and partially corrected in Orb-v3 (7-8 kcal/mol).

**Figure 3.**
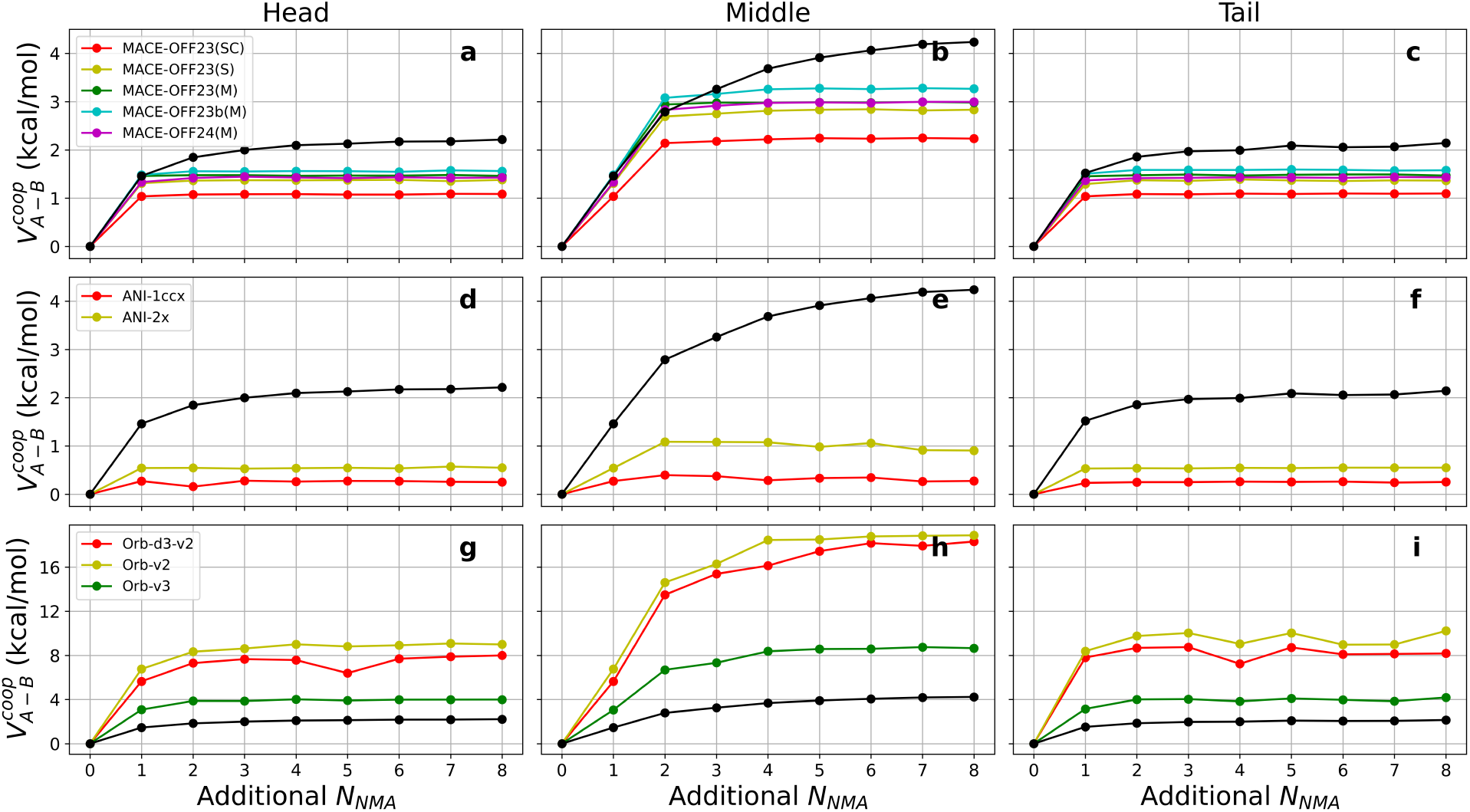
Comparison of H-bond cooperative energies in arc-shaped NMA polymers calculated using QM and MLFFs. The *x*-axis represents the number of NMA units in the polymer, while the *y*-axis shows cooperative energy in kcal/mol. Black dots and lines represent the QM reference, and colored dots and lines correspond to various MLFF predictions as indicated in the legends. (a-c) Cooperative energies for head, middle, and tail H-bonds using MACE-OFF models. Legend labels SC, S, and M denote the soft-core, small, and medium model variants, respectively. (d-f) Cooperative energies for head, middle, and tail H-bonds using ANI models. (g-i) Cooperative energies for head, middle, and tail H-bonds using Orb models.

In ANI and MACE-OFF models, cooperativity reached a plateau immediately after adding a single NMA on either side, with no further increase upon polymer elongation. MACE-OFF models therefore correctly reproduced the cooperative effect of adding one NMA, while underestimating the effect of further elongation. All the MACE-OFF variants performed equivalently in cooperative energy, except for 23-SC, which exhibited a observable drop, likely due to its modified repulsive atomic interactions that are specifically introduced for free energy perturbation calculations [37]. Different from ANI and MACE-OFF, cooperativity in Orb models increased progressively and converged after the addition of five or more NMAs, closely resembling the QM profile.

### 3.3 Cooperativity in presence of water molecules

To further assess the impact of solvent, all polymer chain termini were capped with water molecules (Figure 4, Figure S3 for linear polymers). One water molecule formed a single H-bond with the amide group, while three additional water molecules interacted with the carbonyl group via two H-bonds and formed two additional H-bonds among themselves (Figure 1d, e). In SCS-MP2 reference of NMA dimer H-bond, the cooperative contribution of this four-water flanking arrangement (2.5 kcal/mol, Figure 4a) was comparable to that of two NMA molecules (2.8 kcal/mol, Figure 3b). The MACE-OFF models closely reproduced the contribution, yielding values in the range of 2.2-2.9 kcal/mol. ANI models continued to underestimate the cooperativity, suggesting a limited sensitivity to solvent-induced effects (Figure 4d). In contrast, the Orb models showed smaller deviations in cooperative energy for the water-flanked dimers than for the NMA-flanked ones: Orb_d3-v2 predicted a value of 1.5 kcal/mol, while Orb-v2 yielded 2.8 kcal/mol, both in reasonable agreement with the QM reference (Figure 4g).

**Figure 4.**
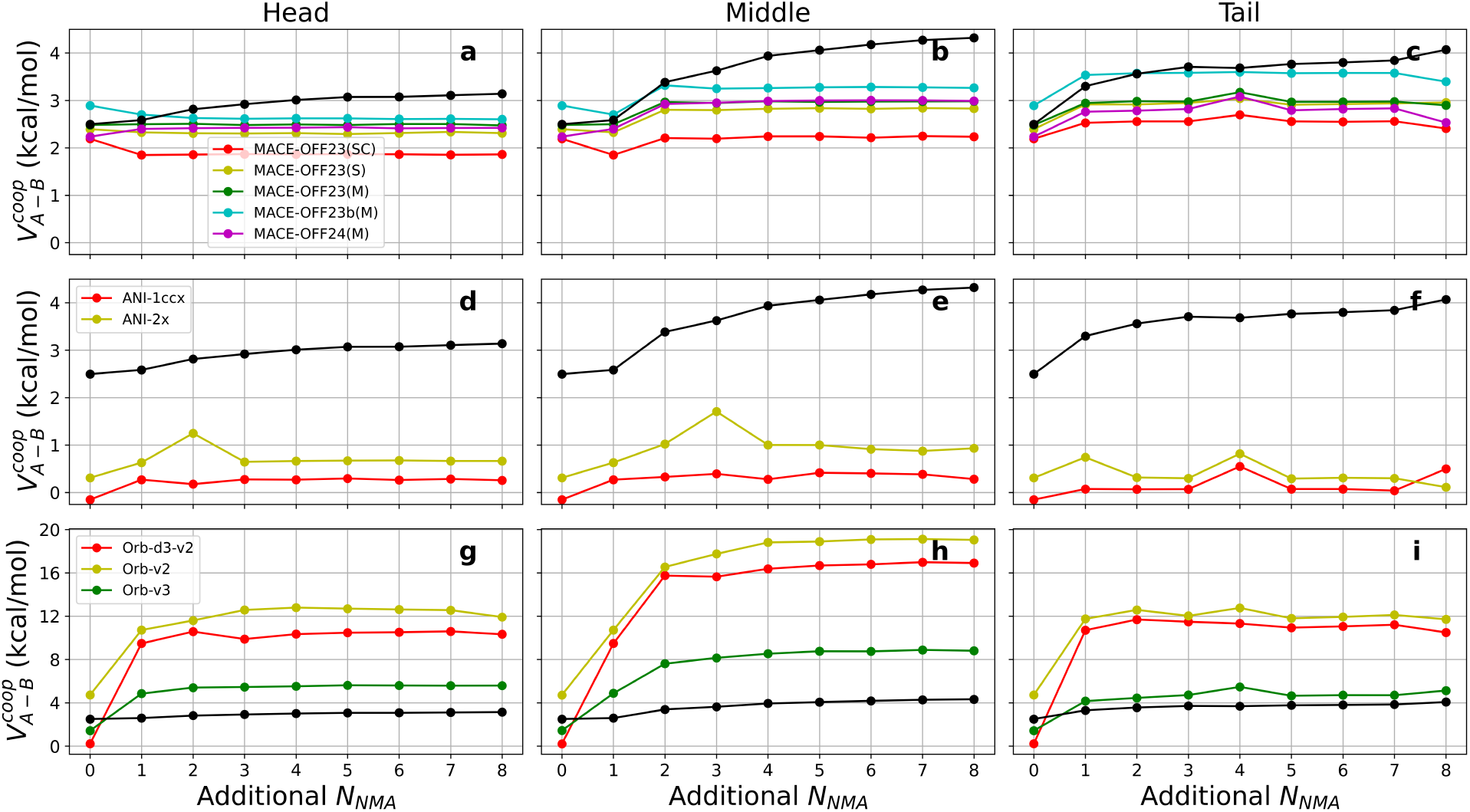
Cooperative energy in arc-shaped NMA polymers in the presence of water molecules. Data organization and representation are identical to those in Figure 3.

Chain extension was performed by inserting additional NMA between the terminal NMA and the adjacent water molecule(s). Due to the contribution of terminal water molecules, the maximum cooperativity observed in SCS-MP2 calculations increased to 3.1 kcal/mol at the head H-bond and 3.9 kcal/mol at the tail H-bond (Figure 4a, c), compared to 2.2 kcal/mol in the NMA-only system. Among the MLFFs, MACE-OFF models closely reproduced the QM reference values, predicting cooperativities of 2.2-2.4 kcal/mol at the head and 2.6-3.5 kcal/mol at the tail. These results suggest that the polarizability of both water and NMA in MACE-OFF aligns well with QM behavior. ANI models still consistently underestimated cooperativity upon elongation, indicating limited response to environmental effects. Orb models continued to overestimate cooperativity, and terminal values exceeded those of the NMA-only system by about 2 kcal/mol, suggesting the Orb water model is reasonably accurate, while the primary source of deviation lies in its NMA representation. Moreover, the trend of increasing cooperativity with NMA extension under the Orb models was qualitatively consistent with QM, which indicates Orb may effectively capture interaction effects of longer range than ANI and MACE-OFF.

In addition to geometric failures observed in the Orb models, outliers in energetics were also identified for MACE-OFF and ANI due to imperfect geometry optimization. Compared to MM force fields, geometry optimization using MLFFs proved more challenging, especially for larger systems. This is consistent with the common assumption that the potential energy surfaces of MLFFs may be less smooth than those of classical FFs. As chain length increased, it became increasingly difficult to locate an energy minimum while maintaining the planarity required for analyzing H-bond cooperativity.

### 3.4 H-bond distance in NMA polymers

The H-bond distance serves as an indicator of interaction strength, with QM-optimized geometries showing progressively shorter bond lengths as cooperativity increases. Among the MLFFs, both ANI and MACE-OFF reproduced dimer H-bond distances in good agreement with QM, indicating their reliability in modeling direct interactions between two NMA molecules (Figure 5, Figure S4 for linear polymers). On the other hand, since the Orb models failed to produce stable H-bonded polymer geometries under standard energy minimization, QM-optimized geometries were directly used to compute the cooperative energies shown in Figure 3. As a result, Orb models were excluded from the analysis of hydrogen bond distance trends in Figure 5.

**Figure 5.**
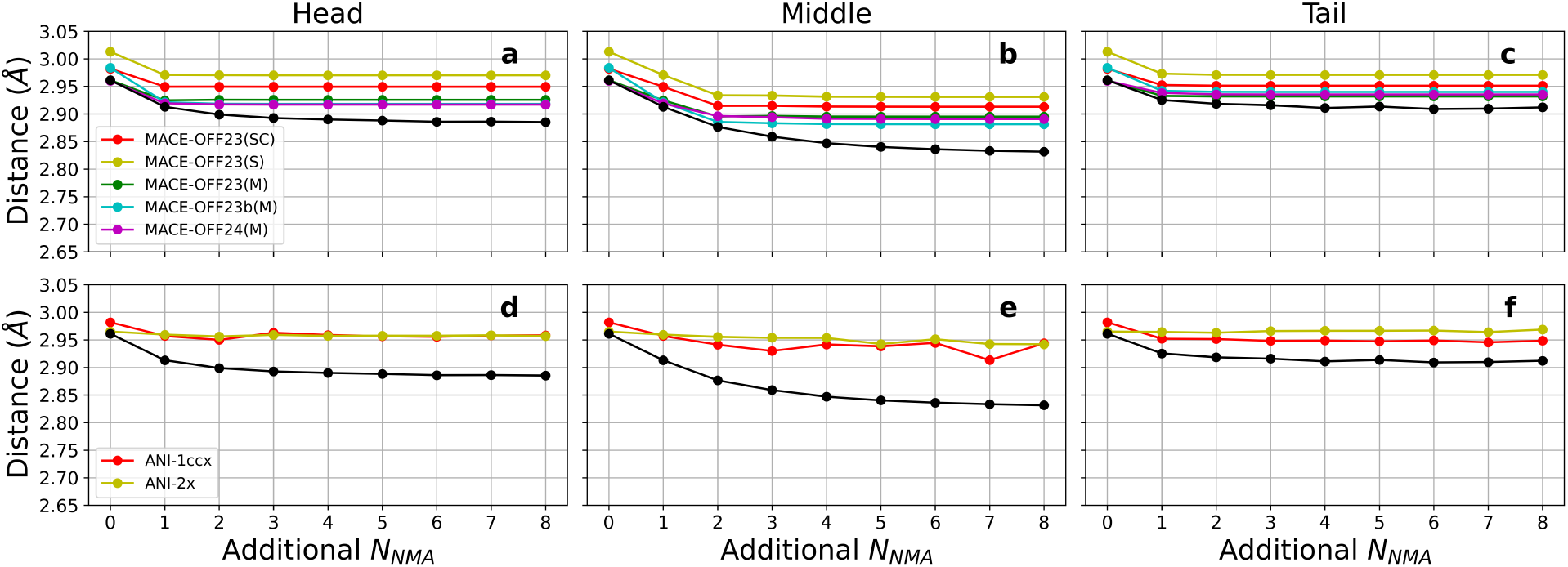
Comparison of H-bond distances in arc-shaped NMA polymers calculated using QM and MLFFs. The *x*-axis represents the number of NMA units in the polymer, while the *y*-axis shows the distance between O_*A*_ and N_*B*_. Other data organization and representation are identical to those in Figure 3.

Upon the addition of one NMA molecule on each side of a dimer, the H-bond distances predicted by MACE-OFF models, including 23-SC, correlated well with their corresponding cooperative energies (Figure 5a, b, c). Distance shortening was evident for both termini, with further reductions in the middle H-bonds of the polymers, consistent with the QM trend. These results also indicate that M-sized MACE-OFF models offer a more accurate description of H-bond distances compared to the S counterpart. Furthermore, as with the energy profiles, the magnitude of distance reduction in MACE-OFF variant was no longer pronounced for further NMA chain extension.

The cooperative effects on H-bond distance in ANI models were much weaker than MACE-OFF and notable deficiencies of optimization in longer polymer were also observed (Figure 5d, e, f). While ANI-1ccx captured a slight distance shortening consistent with cooperative effects, ANI-2x failed to reflect any significant trend. This discrepancy contrasts with their energy profiles, where ANI-2x exhibited more pronounced responses than ANI-1ccx.

In summary, MACE-OFF variants provided the best agreement with QM data for both cooperative energy and H-bond distances. Notably, the polarizability of both NMA and water was correctly captured in MACE-OFF models, reproducing the expected QM trends. Among MACE-OFF variants, M-sized models outperformed S and 23-SC models, but no significant differences were observed between MACE-OFF23 and MACE-OFF24. Although ANI models correctly reproduced NMA dimer H-bonding geometries, they failed to capture the cooperative effects on both energy and H-bond distance. Orb-v2 models were unable to reproduce the correct dimer geometry and drastically overestimated the cooperative energy for NMA, but the polarizability of water was better depicted. Additionally, while MACE-OFF and ANI captured cooperative effects only upon the addition of one NMA, Orb models more closely resembled the QM trend, where cooperative effects continued to accumulate with further NMA chain extension.

## 4 Discussion

Quantifying cooperativity provides a sensitive metric for evaluating a force fields ability to generalize across electrostatic environments and intermolecular interactions. In our previous study,[34] we benchmarked classical MM force fields using the same NMA polymer systems. Additive force fields, by construction, yield strictly zero cooperativity due to the absence of polarization in their potential energy functions. In contrast, polarizable models such as Drude and AMOEBA explicitly incorporate electronic induction and showed clear improvements over additive FFs in reproducing cooperative effects. However, even these advanced FF models underestimated cooperative energy by approximately 20% and 40%, respectively, when compared to QM references.

Leveraging the growing availability of high-level QM data and computational resources, MLFFs offer the potential to model complex systems with improved generality and accuracy. In this study, we assessed three representative MLFFs (ANI, MACE-OFF, and Orb) using the NMA benchmark system designed to probe electrostatic induction and dispersion contributions across increasing chain lengths. Despite differences in architectures and training data, all three models were trained on QM-calculated energies and forces with the goal of achieving broad chemical transferability. A key aspect of comparison lies in their capacity to capture polarizability and long-range interactions — factors central to H-bond cooperativity. As such, the cooperativity trend in NMA polymers provides a stringent test of how well each MLFF models many-body electrostatics across molecular chains.

To explain the performance of long-range cooperativity in MLFFs, we analyze the induced molecular energy changes in more detail. In machine learning potentials, the total energy of a system is expressed as the sum of atomic energies. Assuming molecule *A* contains *m* atoms, *B* contains *n* atoms, and *C* contains *l* atoms, the total energy of the three-body system is given by:

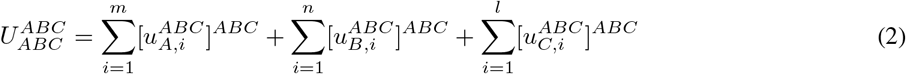

Here, 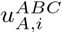 denotes the atomic energy of atom *i* in molecule *A* calculated in the context of the *ABC* complex, and the bracket with superscript *ABC* indicates that the geometry corresponds to the full *ABC* system. This differentiates the notation from the total internal energy 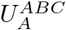 of molecule *A* at the same geometry, as introduced in Eq. 1.

Accordingly, the cooperative energy in Eq. 1 can be rewritten in terms of atomic energy contributions as:

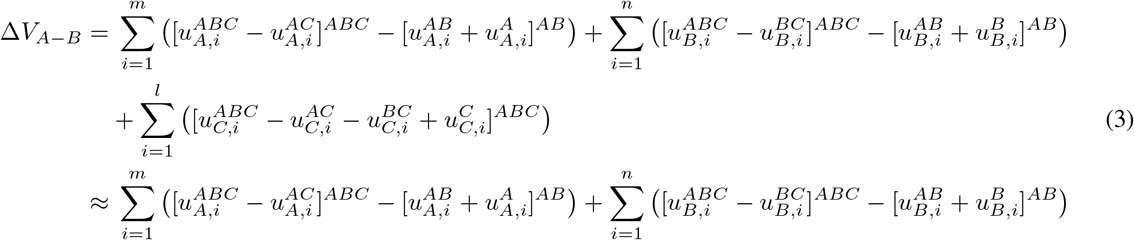

The approximation holds because the third term, 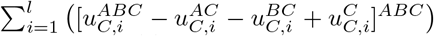, is close to zero. This is because, at an identical geometry, the total energy perceived by molecule *C* from molecules *A* and *B* is nearly equivalent whether considered as a whole (*ABC*) or in pairs (*AC* and *BC*).

To examine the cooperativity change as more NMA units are added, we extend molecule *C* from *N* − 1 to *N* units. The resulting change in cooperative energy is:

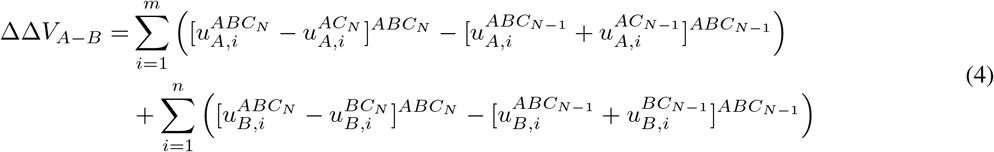

As a case study, we analyze the head H-bond in NMA polymers using the MACE-OFF 23-M model. Eq. 4 can be rearranged into four components:

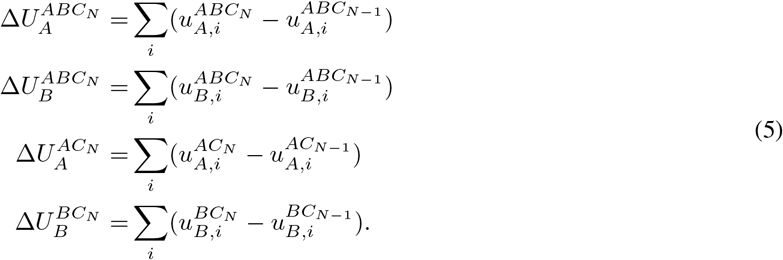

The superscripts were omitted for clarity, because in this case the geometric influence from *C*_*N*_ is small. These terms reflect the influence of the additional NMA in molecule *C* on the energies of *A* and *B*. Figure 6 presents these energy components for *N* = 0, 1, 2, 3, corresponding to NMA dimers, trimer, tetramer, and pentamers, respectively. The results indicate that when *N* = 0, molecule *A* perceives a direct influence from *B*. At *N* = 1, *A* receives minimal influence from *C*, while *B* is affected by both *A* and *C*. When *N* = 2 or 3, the induced effects sharply diminish. Notably, the shortest interatomic distance between molecule *C*_2_ and molecule *B* (or *C*_1_ and *A*) is approximately 6 Å, which lies at the cutoff boundary of the MACE-OFF 23-M model. This suggests that only nearest-neighbor NMAs significantly contribute to cooperative energy, while next-nearest neighbors lie almost beyond the effective interaction range of the model.

**Figure 6.**
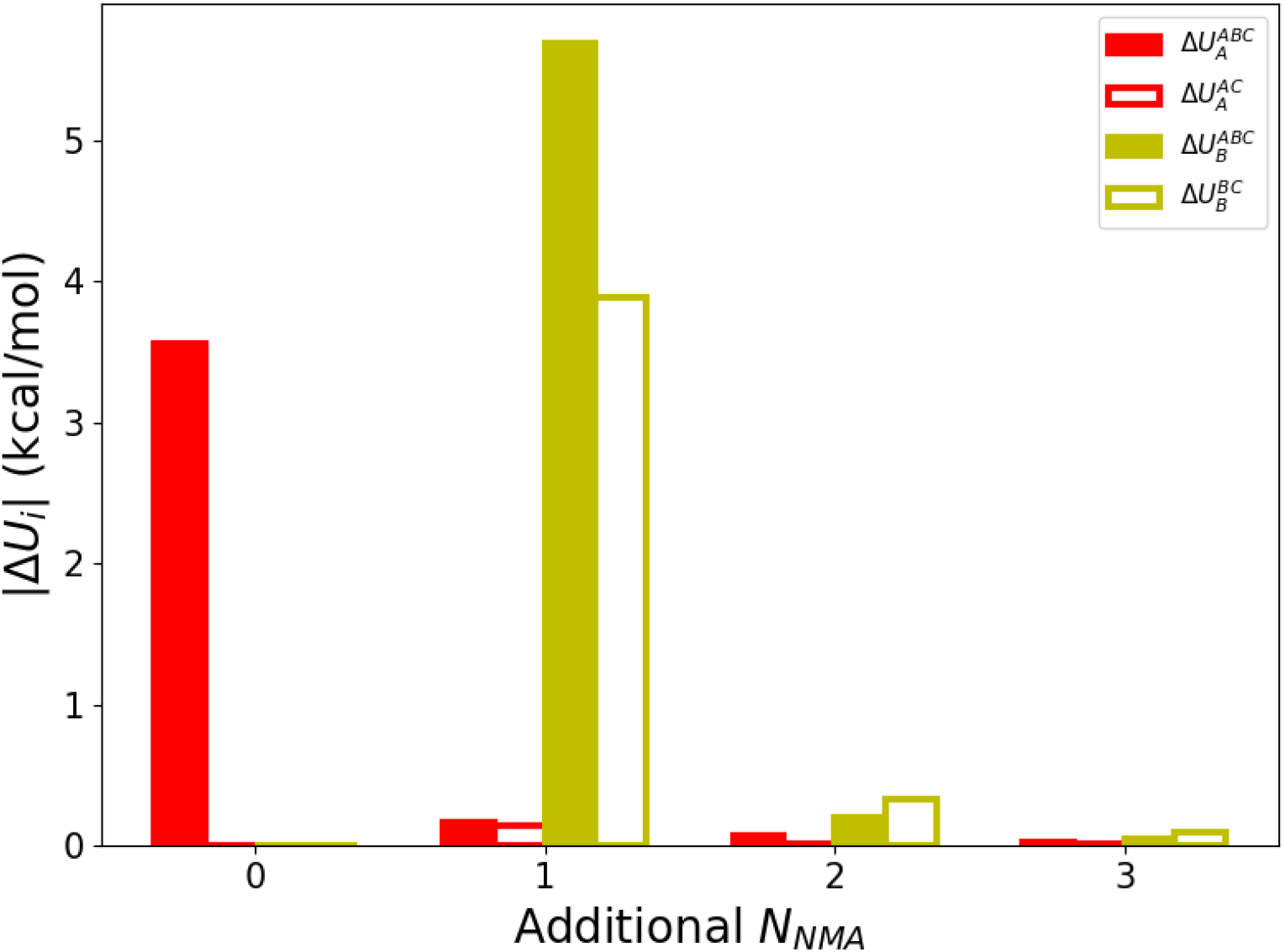
The atomic potential energy differences of A and B molecules relative to the A-B dimer, calculated along the arc polymers of 3,4,5 units using the MACE-OFF 23-M model.

None of the MLFFs fully reproduced the cooperative energy across all chain lengths with quantitative fidelity. Both ANI and MACE-OFF failed to capture the growth of cooperative effects as the number of NMA units exceeded two, highlighting limitations in modeling long-range polarization. ANI, while successful in modeling small molecules, consistently underestimated cooperative energy, reflecting its lack of explicit electrostatic induction. MACE-OFF, by contrast, achieved better agreement at short to medium ranges, likely due to its rotationally equivariant architecture and exposure to peptide-like fragments and solvated molecular complexes during the training. Nonetheless, its sensitivity plateaued beyond trimer systems, suggesting a finite non-bonded perception range.

Orb demonstrated the clearest monotonic rise in cooperativity with chain length, indicating a strong response to long-range correlations. This performance likely arises from its attention-based architecture that incorporate receptive field. However, Orb significantly overestimated the magnitude of cooperative energy, implying an over-polarizable potential.

This may result from its pre-training strategies that prioritize data diversity and generalization but do not sufficiently sample bio-organic environments. While Orb captures the qualitative trend of cooperativity, its quantitative predictions deviates substantially from QM references.

These findings highlight a broader challenge in MLFF development: while modern architectures can encode complex spatial correlations, accurately capturing cooperative effects requires more than architectural sophistication or transfer learning. The large discrepancies between ML models reflect a combination of architectural biases, training data gaps, and the difficulty of learning subtle many-body polarization phenomena from general-purpose datasets.

Our results point to the need for better benchmarks and descriptors of cooperativity. Current MLFF evaluations often focus on local accuracy metrics (e.g., forces, energies, torsion scans), which may overlook systematic deficiencies in long-range interactions. As cooperative H-bonding is ubiquitous in biomolecular systems, we propose that quantifying cooperativity in controlled model systems should become a standard component of MLFF validation. Adopting this approach can help pinpoint force field limitations and inform the development of next-generation neural network potentials that incorporate inductive and many-body effects in a physically meaningful way.

## 5 Conclusion

This study evaluated the capability of MLFFs to reproduce cooperative H-bonding interactions in NMA polymers, a representative system for biomolecular noncovalent interactions. By quantifying the cooperative energy and structural changes, we assessed three state-of-the-art MLFFs — ANI, MACE-OFF, and Orb — against high-level QM calculations. Significant differences were observed among the models, demonstrating that H-bond cooperativity can serve as a critical benchmark for evaluating the accuracy and transferability of MLFFs.

Universal MLFFs have rapidly evolved with the general aim of simulating complex chemical and biological systems. The benchmark results presented in this work underscore the importance of evaluating MLFFs not only in terms of pairwise accuracy but also in their treatment of collective many-body phenomena. H-bond cooperativity provides a rigorous and physically interpretable benchmark for assessing long-range induction and dispersion effects, which are crucial for accurately modeling biomolecules. Our findings suggest that while current MLFFs show promising accuracy for isolated interactions, further development is needed to better account for cooperative behavior across extended systems, particularly in biological and condensed-phase applications.

## Acknowledgements

This work was supported by the Pioneer and Leading Goose R&D Program of Zhejiang (2023C03109, 2024SSYS0036), the National Natural Science Foundation of China (32171247, 21803057). We thank the Westlake University Supercomputer Center for computational resources and related assistance.

## Conflict of interest

No conflict of interest.

## Notes

### Competing Interest Statement

The authors have declared no competing interest.

### Summary of Updates

Add additional analysis on the distance dependence. One new figure is added.

