## Supplementary Materials for "Benchmarking Universal Machine Learning Force Fields with Hydrogen-Bonding Cooperativity"

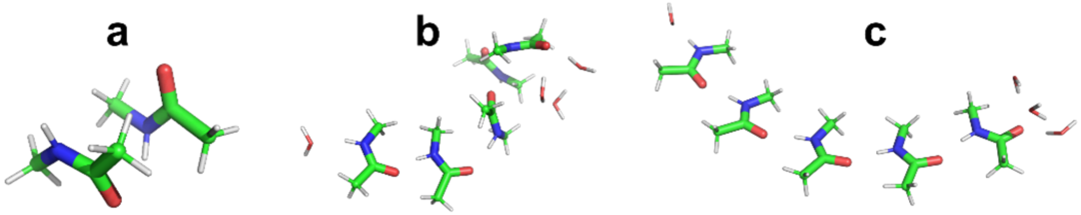


Figure S1. **Examples of Orb optimized NMA polymers.** (a) Orb-v2 optimized dimer without planarity restraints. (b) Optimized pentamer with planarity restraints and a moderate energy tolerance (1.1 kcal/mol/Å). (c) QM optimized pentamer for cooperative energy calculation.

Figure S2-S4 Cooperativity calculated for linear-shaped NMA polymers.

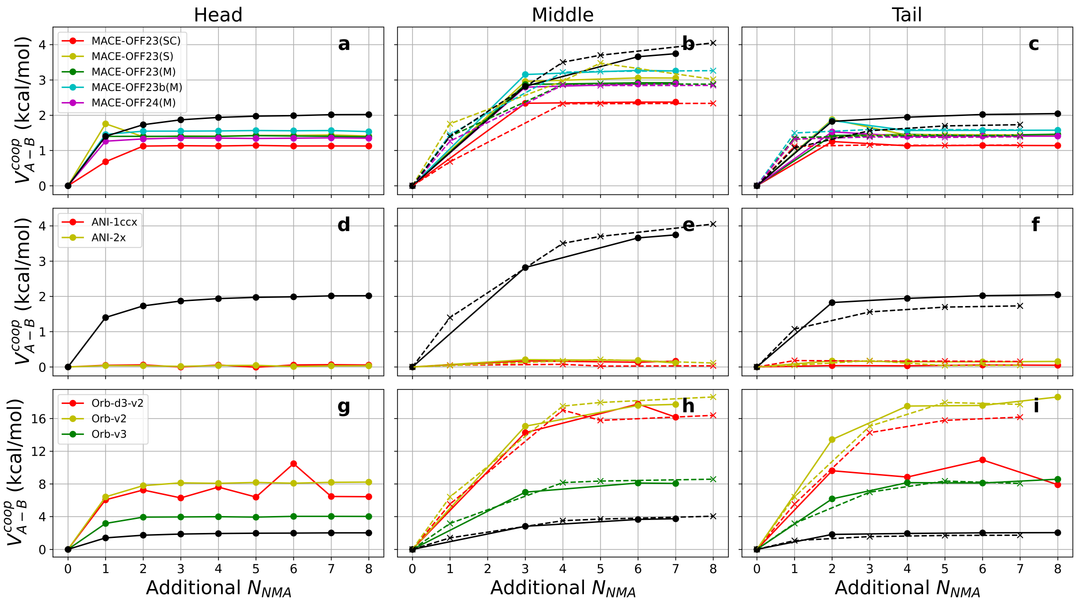


FIGURE S2. **Comparison of H-bond cooperative energies in linear-shaped NMA polymers calculated using QM and MLFFs.** The $x$-axis represents the number of NMA units in the polymer, while the $y$-axis shows cooperative energy in kcal/mol. Dots and lines represent the H-bonds in syn conformation, and cross and dashed-lines represent the H-bonds in anti. Black is for QM reference, and other colors correspond to various MLFF predictions as indicated in the legends. (a-c) Cooperative energies for head, middle, and tail H-bonds using MACE-OFF models. Legend labels SC, S, and M denote the soft-core, small, and medium model variants, respectively. (d-f) Cooperative energies for head, middle, and tail H-bonds using ANI models. (g-i) Cooperative energies for head, middle, and tail H-bonds using Orb models.

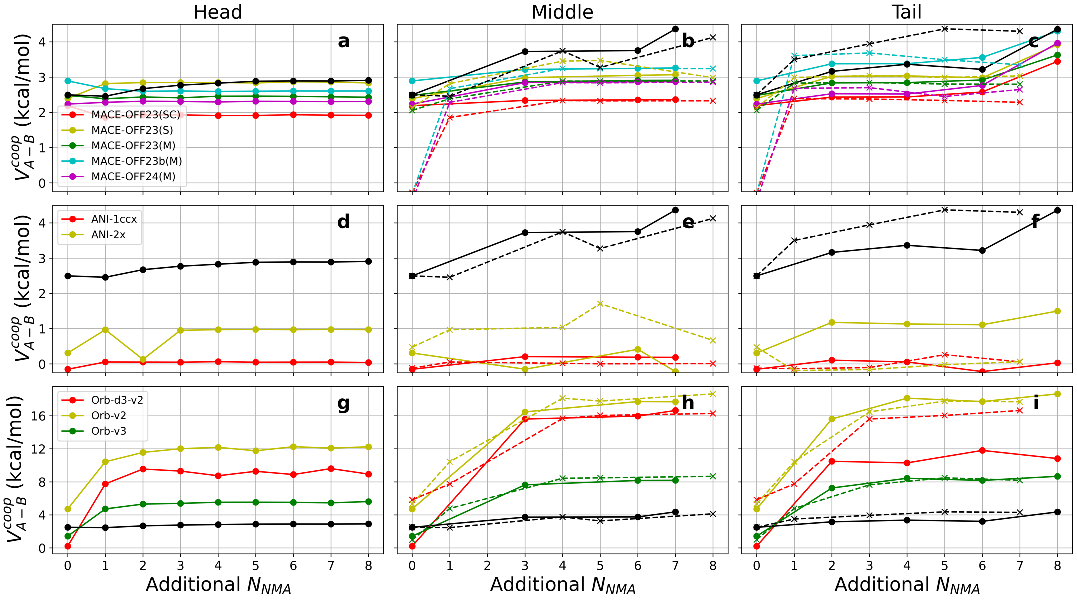


FIGURE S3. **Cooperative energy in linear-shaped NMA polymers in the presence of water molecules.** Data organization and representation are identical to those in Figure S2.

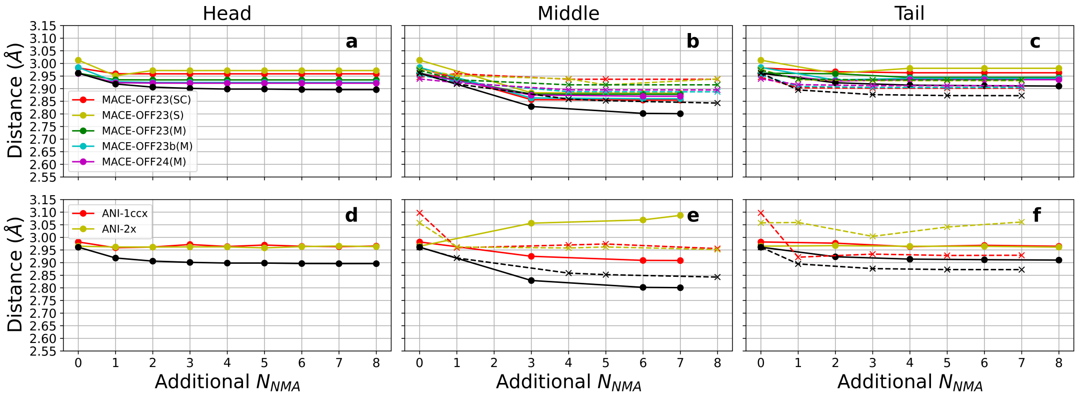


FIGURE S4. **Comparison of H-bond distances in linear-shaped NMA polymers calculated using QM and MLFFs.** The $x$-axis represents the number of NMA units in the polymer, while the $y$-axis shows the distance between $O_A$ and $N_B$. Other data organization and representation are identical to those in Figure S2.

Figure S5-S8 Cooperative energies on QM geometries without energy-minimization using MLFFs.

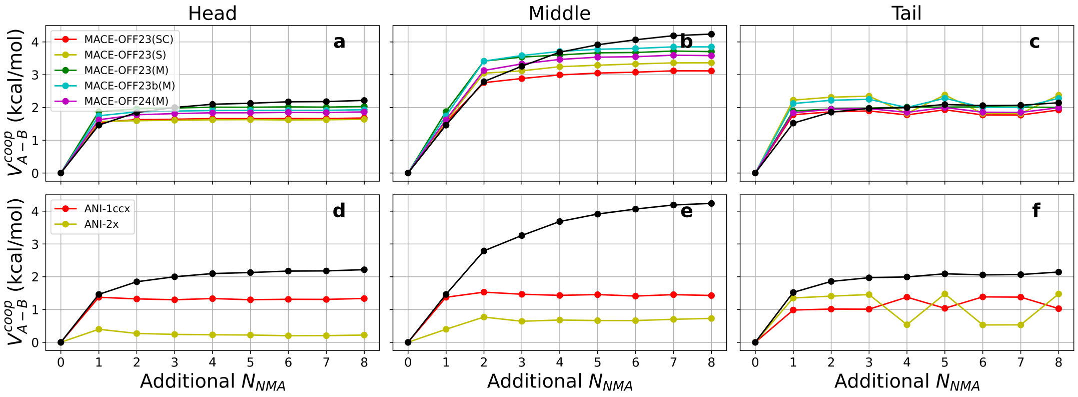


FIGURE S5. **H-bond cooperative energies in arc-shaped NMA polymers calculated using QM optimized geometries.** The $x$-axis represents the number of NMA units in the polymer, while the $y$-axis shows cooperative energy in kcal/mol. Black dots and lines represent the QM reference, and colored dots and lines correspond to various MLFF predictions as indicated in the legends. (a-c) Cooperative energies for head, middle, and tail H-bonds using MACE-OFF models. Legend labels SC, S, and M denote the soft-core, small, and medium model variants, respectively. (d-f) Cooperative energies for head, middle, and tail H-bonds using ANI models. (g-i) Cooperative energies for head, middle, and tail H-bonds using Orb models.

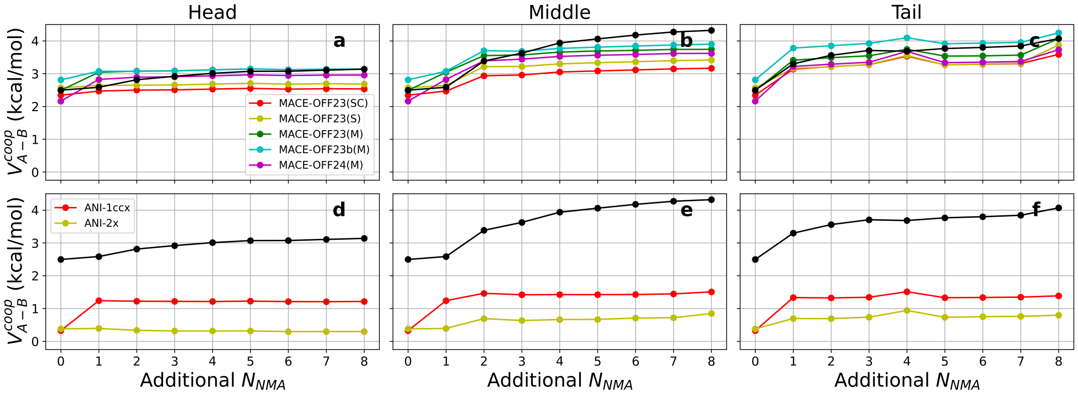


FIGURE S6. **H-bond cooperative energies in arc-shaped NMA polymers in the present of water molecules calculated using QM optimized geometries.** Data organization and representation are identical to those in Figure S5.

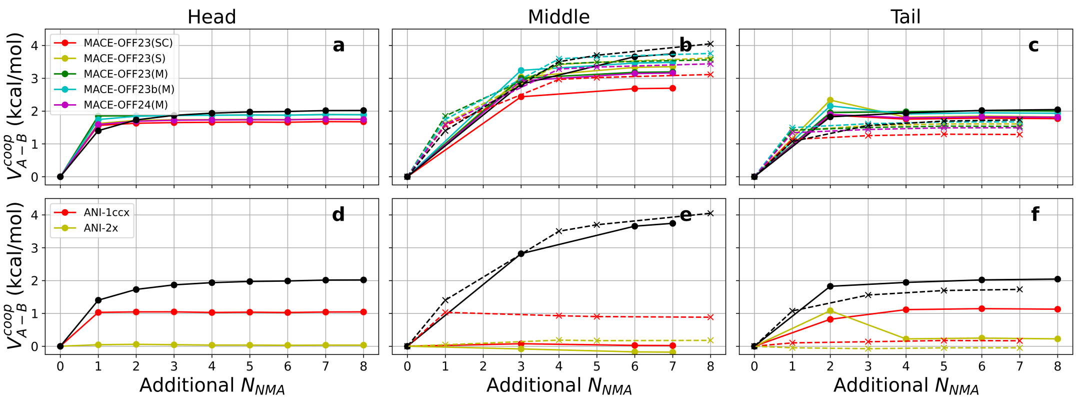


Figure S7. **H-bond cooperative energies in linear-shaped NMA polymers calculated using QM optimized geometries.** Dots and lines represent the H-bonds in syn conformation, and cross and dashed-lines represent the H-bonds in anti. Other data organization and representation are identical to those in Figure S5.

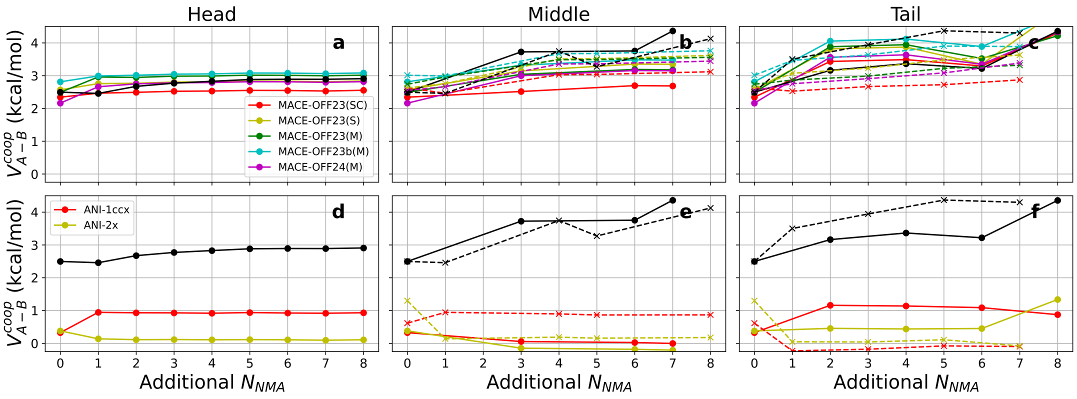


Figure S8. **H-bond cooperative energies in linear-shaped NMA polymers in the present of water molecules calculated using QM optimized geometries.** Data organization and representation are identical to those in Figure S7.

| Variants | Website | Access date |
|---|---|---|
| 23-SC | https://github.com/jharrymoore/MACE-OFF23-SC | 18-Nov-24 |
| 23-SMALL<br>23-MEDIUM<br>23b-MEDIUM<br>24-MEIDUM | https://github.com/ACEsuit/mace-off | 19-Feb-25 |
| ANI-1ccx<br>ANI-2x | https://github.com/aiqm/ani-model-zoo/tree/master/resources | 27-Feb-25 |
| Orb-v2<br>Orb-d3-v2 | https://github.com/orbital-materials/orb-models | 27-Feb-25 |
| Orb-v3 | | 18-Apr-25 |

Table S1. **Access Information of Machine Learning Potentials.**
